# PYPE: A Python pipeline for phenome-wide association (PheWAS) and mendelian randomization in investigator-driven phenotypes and genotypes of biobank data

**DOI:** 10.1101/2022.12.10.519906

**Authors:** Taykhoom Dalal, Chirag J. Patel

## Abstract

**Motivation:** Phenome-wide association studies (PheWASs) serve as a way of documenting the relationship between genotypes and multiple phenotypes, helping to uncover new and unexplored genotype-phenotype associations (known as pleiotropy). Secondly, Mendelian Randomization (MR) can be harnessed to make causal statements about a pair of phenotypes (e.g., does one phenotype cause the other?) by comparing the genetic architecture of the phenotypes in question. Thus, approaches that automate both PheWAS and MR can enhance biobank scale analyses, circumventing the need for multiple bespoke tools for each task by providing a comprehensive, end-to-end pipeline to drive scientific discovery.

**Results:** We present PYPE, a Python pipeline for running, visualizing, and interpreting PheWAS. Our pipeline allows the researcher to input genotype or phenotype files from the UK Biobank (UKBB) and automatically estimate associations between the chosen independent variables and the phenotypes. PYPE also provides a variety of visualization options including Manhattan and volcano plots and can be used to identify nearby genes and functional consequences of the significant associations. PYPE additionally provides the user with the ability to run Mendelian Randomization (MR) under a variety of causal effect modeling scenarios (e.g., Inverse Variance Weighted Regression, Egger Regression, and Weighted Median Estimation) to identify possible causal relationships between phenotypes.

**Availability and Implementation:** PYPE is a free, open-source project developed entirely in Python and can be found at https://github.com/TaykhoomDalal/pype. PYPE is published under the Apache 2.0 license and supporting documentation can be found at the aforementioned link.

## 1 Introduction

While genome-wide association studies have been critical for determining the relationship between genetic variants along the genome and a phenotype, phenome-wide association studies allow us to query the reverse, determining the relationship between phenotypes along the phenome and a genetic variant (Denny et al. 2010). PheWAS have been shown to replicate known GWAS results (Denny et al. 2013) and to improve upon GWASs by helping to identify shared biological mechanisms across phenotypes (pleiotropy), reveal previously unknown associations between phenotypes, and identify novel associations (Diogo et al. 2018; Tyler, Crawford, and Pendergrass 2016). The development of tools that can take advantage of biobank scale data sources for performing PheWAS is thus a prerequisite for interpreting and contextualizing genotype-phenotype associations.

Prior PheWAS software includes R PheWAS (Carroll, Bastarache, and Denny 2014), PyPheWAS (Kerley et al. 2022), and DeepPheWAS (Packer et al. 2022), the latter of which uses Plink2, a popular open-source association analysis toolset (Chang et al. 2015), to accelerate their analyses. Packages also exist for visualizing PheWAS results, such as PheWAS-View (Pendergrass et al. 2012) and PheWeb (Gagliano Taliun et al. 2020). While these tools are useful in aggregate, none of these tools by themselves provide an end-to-end pipeline for running a PheWAS, visualizing the results, and interpreting potential causal effects. To this end, we developed a Python-based implementation of a PheWAS pipeline, including downstream functionality that annotates the significant results with nearby genes and functional relevance according to external variant information databases, and can natively run Mendelian Randomization (MR), a method that uses genetic variants to approximate a randomized control trial to make causal inferences about the relationship between an exposure and an outcome (Ebrahim and Davey Smith 2008). While packages for MR exist, such as MendelianRandomization (Yavorska and Burgess 2017) and TwoSampleMR (Hemani, Tilling, and Davey Smith 2017), none of these are incorporated into any previous PheWAS workflow, even though it is a very typical analysis to run along with the PheWAS. To this end, we present PYPE, a computational pipeline, optimized for UKBB data, that is designed to accelerate PheWASs at every step of the process.

## 2. Methods

PYPE facilitates the three main components of a typical PheWAS analysis. First, the user inputs a file containing the independent variables (which may be a set of genotype or phenotype files), a file containing the dependent variables (the set of phenotypes provided by the UKBB), and any other optional arguments (e.g. covariates) and runs the PheWAS analysis. Next, the user can choose to visualize the PheWAS results in a variety of plot types, highlighting significant associations that were found in the PheWAS. Finally, the user can explore the functional consequences of the significantly associated variants and the genes these variants are physically close to, as well as run MR analyses to uncover possible causal relationships for the phenotypes of interest.

### 2.1 PheWAS

The UKBB is a large-scale medical database that provides in-depth genetic, health, and lifestyle data for around half a million UK residents, allowing for a wide variety of exploratory analyses that facilitate biological discovery (Sudlow et al. 2015). With this in mind, we decided to focus our initial pipeline compatibility with this widely used resource to provide the most useful tool for most researchers. With the data provided by the UKBB, users can input either genotype or phenotype data as the independent variables, and phenotype data as the dependent variables in the PheWAS. Users can also specify covariates such as age, sex, genetic principal components, or other phenotypes provided by UKBB for use in the PheWAS. Furthermore, the pipeline scrapes the UKBB website to annotate the specified phenotypes with description and categorization information, allowing users to specify broad classes of phenotypes directly using a link from the UKBB showcase.

PYPE currently uses the default linear regression model from statsmodels in Python to run the mass multivariate regressions, with phenotypes as the dependent variables and genotypes (or phenotypes) as the independent variables, outputting p-values, beta coefficients, and standard errors. Here, the beta coefficients that are produced for each variant-phenotype association indicate whether having the variant increases or decreases the phenotype’s value, a fact that can be used to interpret the biological impact of the variant on the target phenotype. PYPE also supports parallelization using Python’s multiprocessing package. To correct for the occurrence of false positives, users can also specify multiple testing correction methods including Bonferroni correction, false discovery rate correction, Sidak correction, or simple p-value correction methods. For each of these multiple testing correction methods, the total number of association tests run is used as an estimation for the space of tests.

### 2.2 Visualization

PYPE also provides a script to generate a variety of visualizations once the PheWAS has been run. Users can choose between generating traditional Manhattan plots, category enrichment plots, and volcano plots. When generating Manhattan plots, users can specify custom groupings for the phenotypes or can use the categories provided by UKBB, can include annotations for the significant associations, alter plot features such as color maps and transparency, and can plot aggregate-level Manhattan plots (Figure 1) or individual categories with significant associations (Figure 3). Users can also generate category enrichment plots as in Figure 2, plotting the enrichment of significant associations per category in a barplot. Finally, users can generate volcano plots for each independent variable, displaying the significant associations between the variable and the dependent phenotypes, with the ability to annotate the most significant associations as well (Supplementary Note Figure 1).

**Figure 1.**
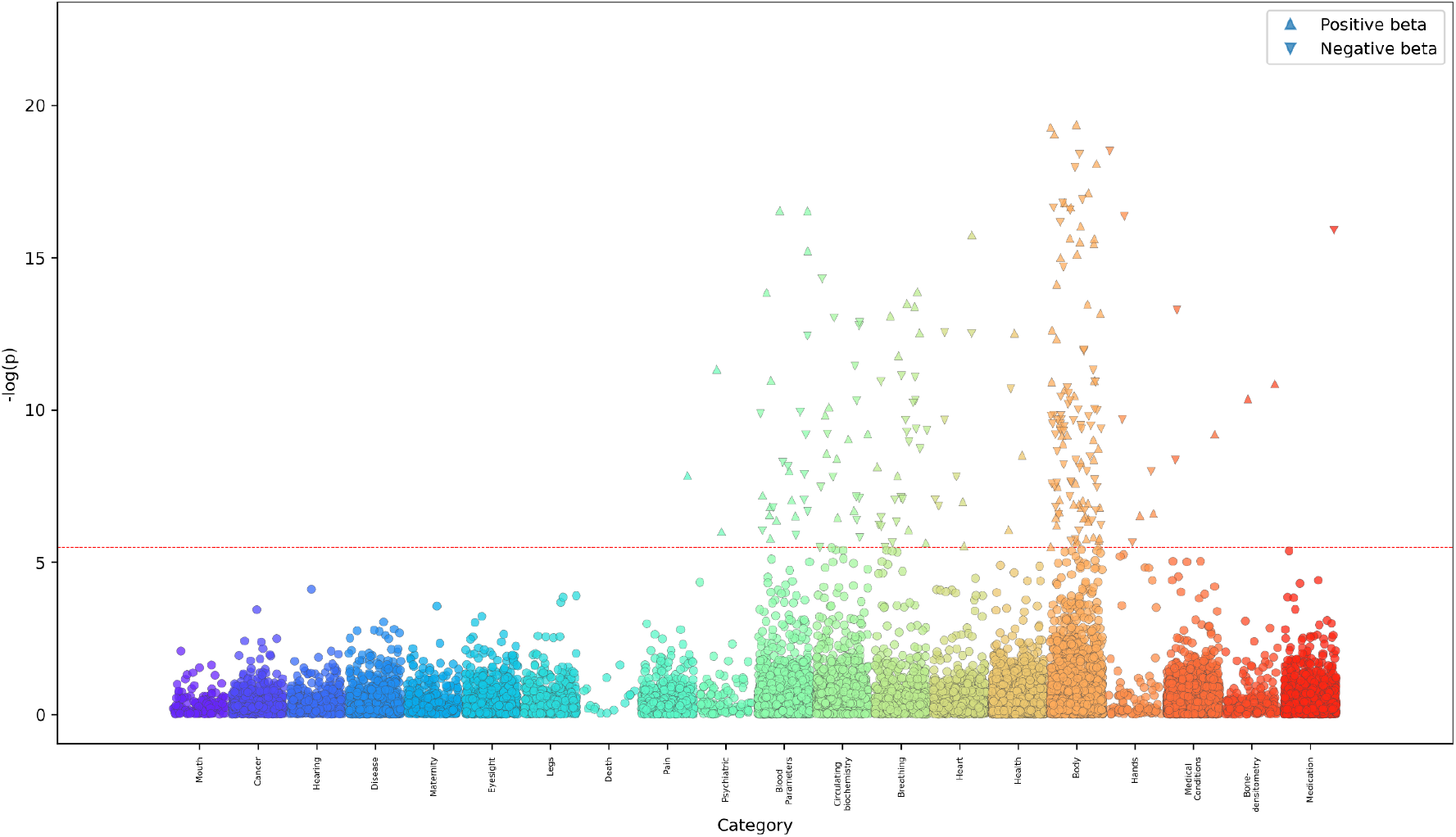
Manhattan plot of aggregate PheWAS results across abdomen, liver, and pancreas age

**Figure 2.**
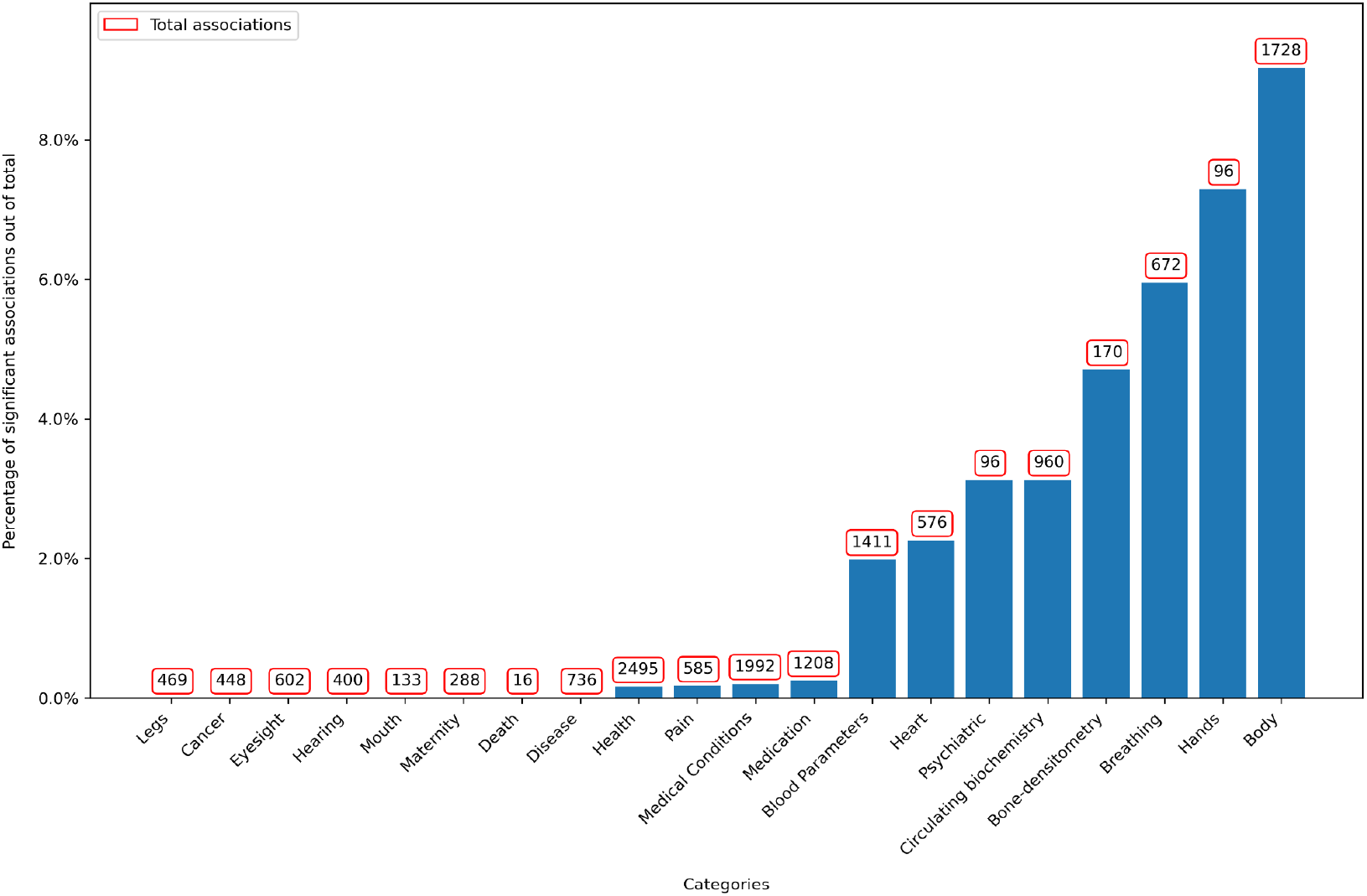
Percentage of each category in terms of the number of significant associations. The number of total associations found is labeled above each category.

**Figure 3.**
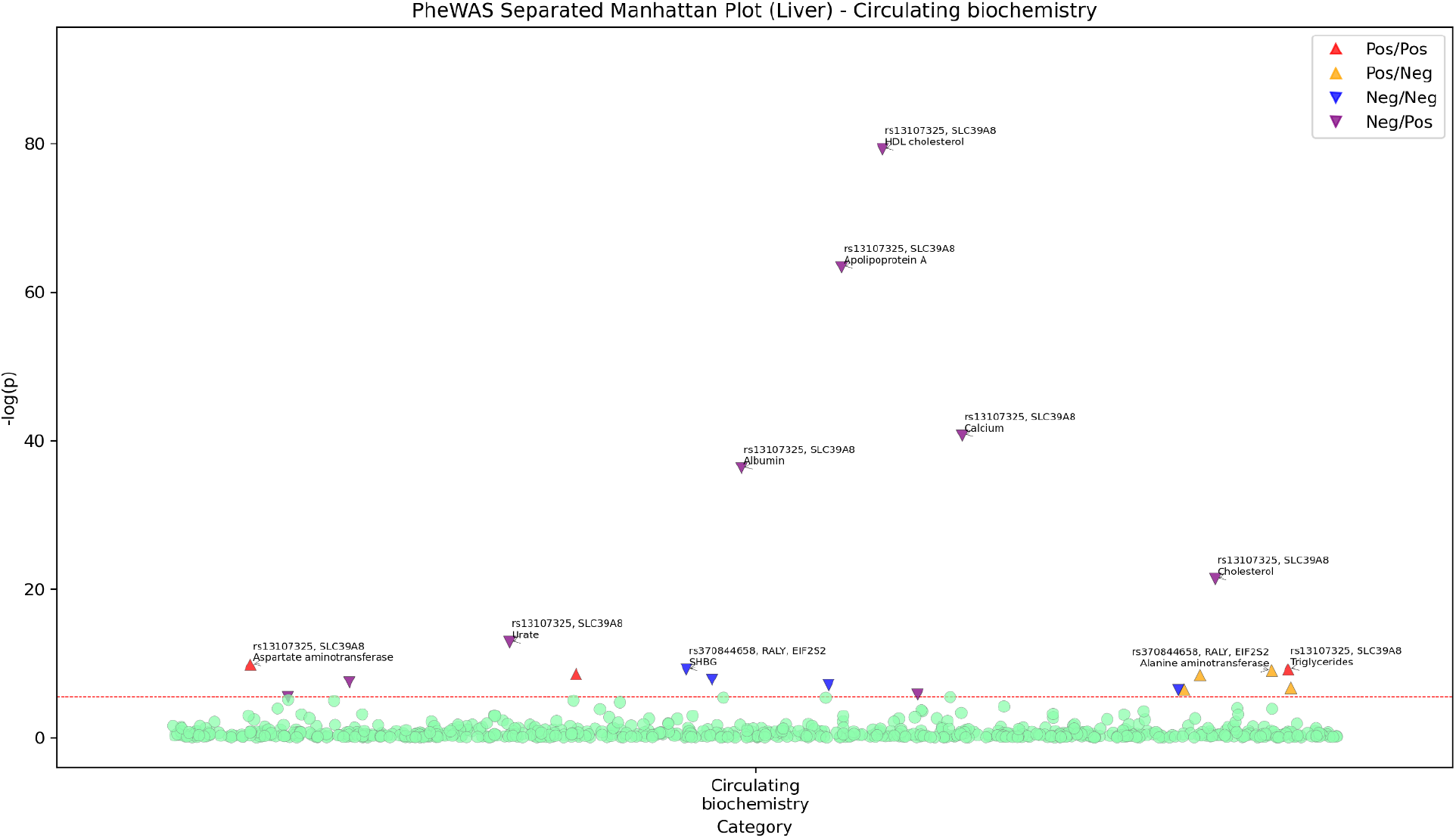
Associations between “Liver aging” variants and circulating biochemistry biomarkers. Note that the directionality of the arrow indicates the sign of the effect size of the PheWAS association, and the color indicates the sign of the effect size of the variant’s association with the aging phenotype defined in Le Goallec et al. 2022. There, a negative effect size indicated accelerated aging and a positive effect size indicated decelerated aging.

### 2.3 Downstream Analysis

If inputting genotype variables, then PYPE provides downstream annotations for the significant associations that are found in the PheWAS. Specifically, PYPE provides three main downstream functions. First, if the user specifies a gene file (which contains information about the location of genes on each chromosome) and the number of kilobases downstream and upstream to look for genes, PYPE can be used to annotate each variant with which genes are close to it on the chromosome. Secondly, users can choose whether to generate summary files for the significant variants using MyVariantInfo and MyGeneInfo (RESTful API) queries to a variety of databases that contain functional variant annotations and their clinical significance (such as ClinVar, dbSNP, CADD, etc) for information about the given variant and its associated genes (Lelong et al. 2022). PYPE will then use this information to create files for each unique, significant variant, describing its functional consequence and the function of the genes it is close to (if this information exists). Lastly, the user can choose to run a two-sample Mendelian Randomization using the genotype data against a second dataset of GWAS results that is either user-specified, or if no dataset is provided, then it is queried from the Open GWAS Project using their RESTful API (Pierce and Burgess 2013; Elsworth et al. 2020). Similar to the TwoSampleMR package, against which our code’s accuracy was verified, the user has a choice between a variety of MR methods, including Inverse Variance Weighted Regression, Egger Regression, and Weighted Median Estimation, providing the ability to assess causal relationships as well as identify bias due to pleiotropic effects (Bowden et al. 2016). Here, the code is a port of the established TwoSampleMR R package for Python.

## Results

To demonstrate the functionality of this pipeline, we applied PYPE on the results of abdomen biological age predictors (Goallec et al. 2022), exploring the genetic architecture of these predictors with cardiometabolic diseases such as type 2 diabetes (T2D) on an out-of-sample dataset from the UKBB. Using the 16 genetic variants implicated in accelerated abdomen, liver, and pancreas aging, we ran the PheWAS against several broad classes of phenotypes found in the UKBB, including physical lab measurements, self-reported medical conditions, linked health outcomes, blood biochemistry, blood count (parameters), and infectious disease antigens. These classes were then manually split into 20 categories based on their labeling in the UKBB, providing a novel classification of the widely used phenotype data fields in the UKBB, a useful resource that can be used going forward in future PheWAS (see Supplementary Note Table 1). We ran the PheWAS with sex, age, ethnicity, and all 40 genetic principal components as covariates, generating 15,000 total associations and 297 significant associations (P < 3.32e-06 Bonferroni corrected with original alpha = 0.05). Finally, we ran MR to assess the possible causal relationship between accelerated abdomen, liver, and pancreas aging and other phenotypes that have been linked to cardiometabolic diseases.

Figure 2 highlights the percentage of significant associations per category, illustrating which phenotypic categories have the greatest percentage of significant associations. We observe the greatest number of significant associations in the Body category, a category that was defined to encapsulate many of the physical lab measures that have to do with adiposity, height, and weight, phenotypes. Furthermore, other top categories include Breathing and Circulating Biochemistry, where Circulating Biochemistry chiefly contains information about cardiometabolic biomarkers such as alanine aminotransferase (ALT), aspartate aminotransferase (AST), gamma-glutamyl transferase (GGT), etc (Chen et al. 2017). For this demonstration, we will focus on the Circulating Biochemistry category results for liver aging, as many of the mentioned enzymes are liver-specific.

Out of the various associations in Figure 3, we observed significant associations for HDL Cholesterol (HDL-C), AST, and ALT. Here, the association between variant rs13107325 and HDL-C indicates that decelerated aging is associated with lower HDL-C. Using the downstream functionality of PYPE, where the base-pair position of the variant on the chromosome is used to map variants to nearby genes (based on user-specified distances), we see that the associated gene for this variant is *SLC39A8*. Another significant association was found between AST and variant rs13107325, with the PheWAS results indicating that with decelerated aging, AST levels are higher. Lastly, the PheWAS found a significant association between the variant rs370844658 and ALT, suggesting that with accelerated aging, ALT levels increase. Using the variant-gene mapping functionality, we find that the associated genes for this variant are *RALY* and *EIF2S2*.

**Table 1.** Mendelian randomization (IVW) results with the biological age variants from Le Goallec et al. 2022 and GWAS results from the UKBB. The red colors indicate significant associations (uncorrected for multiple hypotheses with P < 0.05)

| Predictor | Phenotype | P-value | Coefficient | Standard Error |
| --- | --- | --- | --- | --- |
| **Abdomen** | Hemoglobin A1C | 0.385 | -0.048 | 0.056 |
|  | Body Mass Index | 0.910 | -0.004 | 0.039 |
|  | Glucose | 0.241 | 0.013 | 0.011 |
|  | **Waist Circumference** | **0.000** | **-0.380** | **0.098** |
|  | Waist-Hip Ratio | 0.229 | -0.001 | 0.001 |
| **Liver** | Hemoglobin A1C | 0.634 | -0.031 | 0.066 |
|  | Body Mass Index | 0.817 | 0.023 | 0.097 |
|  | Glucose | 0.961 | 0.001 | 0.011 |
|  | Waist Circumference | 0.141 | -0.232 | 0.158 |
|  | Waist-Hip Ratio | 0.280 | -0.001 | 0.001 |
| **Pancreas** | **Hemoglobin A1C** | **0.020** | **0.159** | **0.069** |
|  | **BMI** | **0.001** | **-0.161** | **0.048** |
|  | **Glucose** | **0.001** | **0.046** | **0.014** |
|  | **Waist Circumference** | **0.000** | **-0.699** | **0.121** |
|  | **Waist Hip Ratio** | **0.000** | **-0.004** | **0.001** |

After running the PheWAS, annotating the significant results with nearby genes, and visualizing the results in various plot types, we ran Mendelian Randomization for the abdomen, liver, and pancreas variants against GWAS significant associations (p < 5e-8) for a variety of phenotypes. IVW Regression was used here to estimate the causal effect of the various age predictors and phenotypes that have known associations with cardiometabolic diseases, finding associations (p < 0.05) between the pancreas aging variants and A1C, BMI, glucose, waist circumference, and waist hip-ratio, as well as between abdomen aging variants and waist circumference.

## Discussion

We present PYPE as an easy-to-use, customizable, and feature-rich tool for running PheWAS studies from start to finish. PYPE allows researchers to specify the options they want at all stages of the analysis pipeline, from execution, to visualization, to any downstream analysis. With this tool, we can abstract away many portions of a typical PheWAS study, allowing more time to be spent on validating and exploring the consequences of the results generated, in contrast to the aforementioned tools (Packer et al. 2022). However, there are some limitations to the current implementation. Chiefly, the tool is currently optimized for the UKBB dataset, and thus many of the functionalities of the tool are not well suited for other datasets. Furthermore, the visualization capabilities are currently limited to static generation of the plots, and the user has to re-generate the results when they want to change an attribute of the plot. However, with the availability of tools such as PheWEB, the latter issue is less of a problem, as PYPE output can be adapted to support interoperability with this tool as well as PheWAS-View, giving the user a variety of visualization choices based on their use case. Lastly, the issue of multiple testing correction of the MR associations is left unaddressed in PYPE, with current literature (Burgess et al. 2019) indicating that conservative approaches to multiple testing correction may be too strict, largely due to the low power of these studies and the fact that the relationships under investigation typically have prior biological support. We acknowledge that PYPE is agnostic to biological claims, and therefore, we recommend associations are reported with false discovery and/or family-wise error rate control. Future versions of PYPE will address these limitations as well as include more visualization options, faster execution, and more varied result annotations, facilitating greater customization for the end user. Tools such as PYPE are essential for reducing the time researchers spend on the data analysis stage and increasing the focus on the result interpretation, a critical stage in the workflow for PheWASs as it is often difficult to differentiate between spurious and true associations.

## Acknowledgements

We would like to thank Harvard Medical School research computing group for access and utilization of the O2 cluster. We also want to acknowledge UK Biobank for providing us with access to the data they collected under project number 52887.

## Author Contributions

T.D. and C.J.P. designed the study. T.D. was involved in creating software, literature search, and wrote the first version of the manuscript. C.J.P. wrote and revised the manuscript.

