## Supplemental Data for "PYPE: A Python pipeline for phenome-wide association (PheWAS) and mendelian randomization in investigator-driven phenotypes and genotypes of biobank data"

### Supplementary Note:

| Old Categories | New Categories |
|---|---|
| Medication | Medication |
| Pain | Pain |
| Eyesight | Eyesight |
| Mouth | Mouth |
| General health | Health |
| Operations | Health |
| Medical conditions | Medical Conditions |
| Medications | Medication |
| Hearing | Hearing |
| Breathing | Breathing |
| Claudication and peripheral artery disease | Legs |
| Cancer screening | Cancer |
| Chest pain | Pain |
| Spirometry | Breathing |
| Blood pressure | Heart |
| Body size measures | Body |
| Body composition by impedance | Body |
| Arterial stiffness | Heart |
| Bone-densitometry of heel | Bone-densitometry |
| Hand grip strength | Hands |
| Acceleration averages | Health |
| Myocardial infarction outcomes | Health |
| Summary Psychiatric | Psychiatric |
| Cancer register | Cancer |
| Summary Administration | Health |
| Summary Maternity | Maternity |

| | |
|---|---|
| Stroke outcomes | Medical Conditions |
| Death register | Death |
| Infectious Disease Antigens | Disease |
| Blood biochemistry | Circulating biochemistry |
| Blood count | Blood Parameters |

Supplementary Table 1. Mapping of categories from the UKBB to custom mappings

| Description | rsID | Samples | p-val | beta | std_error | Gene |
|---|---|---|---|---|---|---|
| HDL cholesterol | rs13107325 | 369277/461744 | 5.75E-80 | -0.02928 | 0.001546 | SLC39A8 |
| Apolipoprotein A | rs13107325 | 367205/461744 | 4.59E-64 | -0.01874 | 0.001109 | SLC39A8 |
| Calcium | rs13107325 | 369307/461744 | 2.10E-41 | -0.00557 | 0.000413 | SLC39A8 |
| Albumin | rs13107325 | 369448/461744 | 5.15E-37 | -0.14498 | 0.011405 | SLC39A8 |
| Cholesterol | rs13107325 | 403306/461744 | 4.19E-22 | -0.04633 | 0.004793 | SLC39A8 |
| Urate | rs13107325 | 402822/461744 | 1.34E-13 | -2.16184 | 0.292041 | SLC39A8 |
| Aspartate aminotransferase | rs13107325 | 401791/461744 | 1.52E-10 | 0.28652 | 0.044744 | SLC39A8 |
| Triglycerides | rs13107325 | 402990/461744 | 6.25E-10 | 0.026443 | 0.004276 | SLC39A8 |
| SHBG | rs370844658 | 365796/461744 | 6.25E-10 | -0.38501 | 0.062258 | RALY, EIF2S2 |
| Alanine aminotransferase | rs370844658 | 403156/461744 | 9.15E-10 | 0.195605 | 0.031942 | RALY, EIF2S2 |
| IGF-1 | rs13107325 | 401115/461744 | 2.72E-09 | 0.138977 | 0.023367 | SLC39A8 |
| Urate | rs370844658 | 402822/461744 | 4.02E-09 | 0.939764 | 0.159724 | RALY, EIF2S2 |
| HDL cholesterol | rs370844658 | 369277/461744 | 1.64E-08 | -0.00478 | 0.000846 | RALY, EIF2S2 |
| LDL direct | rs13107325 | 402565/461744 | 3.42E-08 | -0.02039 | 0.003694 | SLC39A8 |
| Cholesterol | rs1797874 | 403306/461744 | 8.03E-08 | -0.01372 | 0.002556 | TSEN2 |
| Total protein | rs1797874 | 369036/461744 | 2.01E-07 | 0.049531 | 0.009528 | TSEN2 |
| IGF-1 | rs3791675 | 401115/461744 | 3.40E-07 | 0.074109 | 0.014531 | EFEMP1 |
| HDL cholesterol | rs1797874 | 369277/461744 | 4.20E-07 | -0.00417 | 0.000825 | TSEN2 |
| Urea | rs13107325 | 403034/461744 | 1.56E-06 | -0.02758 | 0.005741 | SLC39A8 |
| Total protein | rs13107325 | 369036/461744 | 3.29E-06 | -0.08311 | 0.017866 | SLC39A8 |

Supplementary Table 2. Circulating biochemistry phenotypes significantly associated with Liver aging ($P < 3.32e-06$ using Bonferroni correction with alpha = 0.05)

| rsID | CHR | POS | Non_Effect | Effect | BETA | P | SE |
|---|---|---|---|---|---|---|---|
| rs552571374 | 2 | 25148623 | G | C | -0.202 | 3.90E-08 | 0.037 |
| rs201407787 | 2 | 56071109 | C | T | 0.225 | 3.90E-11 | 0.034 |
| rs3791675 | 2 | 56111309 | C | T | -0.16 | 4.70E-09 | 0.027 |
| rs1797874 | 3 | 12529592 | C | A | -0.14 | 2.10E-09 | 0.023 |
| rs13107325 | 4 | 1.03E+08 | C | T | 0.271 | 1.80E-09 | 0.045 |
| rs12539772 | 7 | 1.21E+08 | T | A | -0.141 | 4.20E-08 | 0.026 |
| rs11111209 | 12 | 1.03E+08 | T | C | 0.201 | 1.80E-08 | 0.036 |
| rs77353655 | 12 | 1.03E+08 | A | G | 0.201 | 4.90E-08 | 0.037 |
| rs76652635 | 13 | 74689496 | A | G | -0.257 | 1.50E-08 | 0.045 |
| rs45515493 | 14 | 21572642 | C | G | -0.195 | 5.10E-09 | 0.033 |
| rs370844658 | 20 | 32679575 | A | ATT | -0.133 | 2.60E-08 | 0.024 |

Supplementary Table 3. The “Liver aging” variants used as a part of the PheWAS.

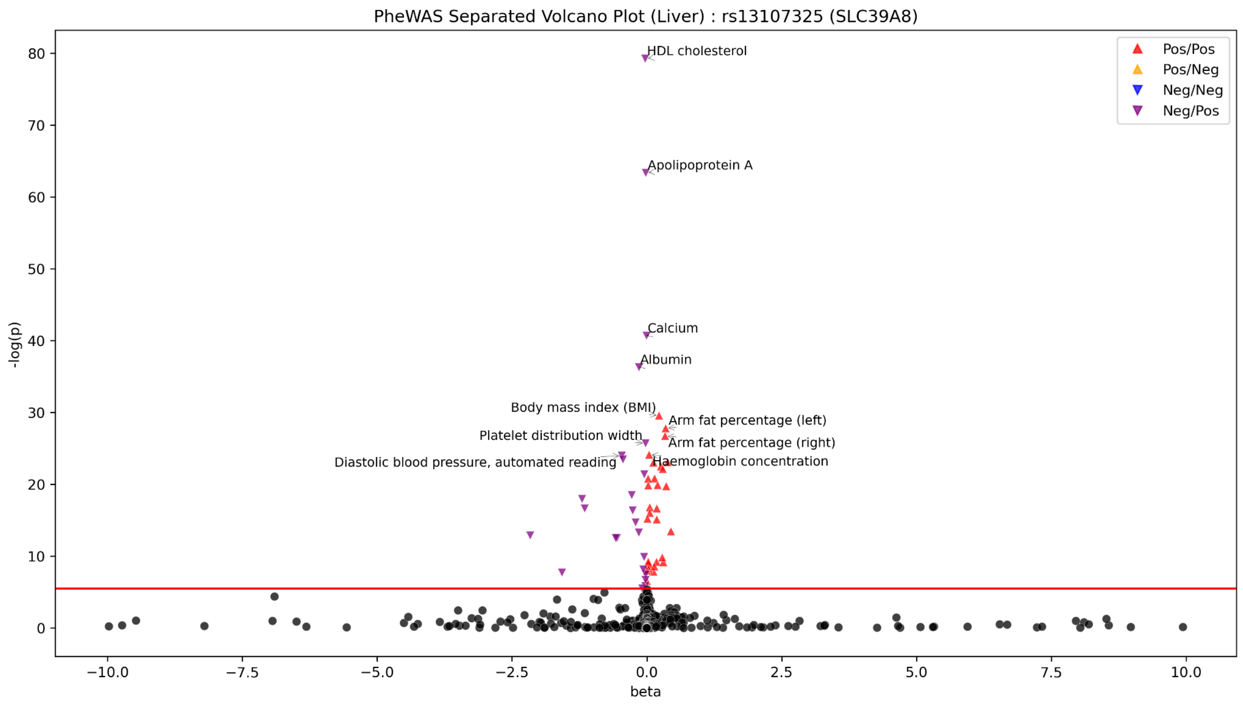


Figure 1. Significant associations between variant rs13107325 and the various phenotypes ran against in the PheWAS. The meaning of the arrow direction and color is identical to in Figure 3, where the arrow indicates the sign of the effect size of the association, and the color indicates the sign of the effect size of the variant’s association with the aging phenotype defined in Le Goallec et al. 2022.
